# Expressing Biologically Active Membrane Proteins in a Cell-Free Transcription-Translation Platform

**DOI:** 10.1101/104455

**Authors:** Shaobin Guo, Amit Vaish, Qing Chen, Richard M. Murray

## Abstract

Cell-free transcription-translation platforms have been widely utilized to express soluble proteins in basic synthetic biological circuit prototyping. From a synthetic biology point of view, it is critical to express membrane proteins in cell-free transcription-translation systems, and use them directly in biocircuits, considering the fact that histidine kinases, G-protein coupled receptors (GPCRs) and other important biosensors are all membrane proteins. Previous studies have expressed membrane proteins in cell-free systems with the help of detergents, liposomes or nanodiscs, but have not demonstrated the ability to prototype circuit behavior for the purpose of testing more complex circuit functions involving membrane-bound proteins. Built on previous efforts, in this work we demonstrated that we could co-translationally express solubilized and active membrane proteins in our cell-free TX-TL platform with membrane-like materials. We first tested the expression of several constructs with β1 and β2 adrenergic receptors in TX-TL and observed significant insoluble membrane protein production. The addition of nanodiscs to the cell free expression system enabled solubilization of membrane proteins. Nanodisc is lipoprotein-based membrane-like material. The activity of β2 adrenergic receptor was tested with both fluorescence and Surface Plasmon Resonance (SPR) binding assays by monitoring the specific binding response of small-molecule binders, carazolol and norepinephrine. Our results suggest that it is promising to use cell-free expression systems to prototype synthetic biocircuits involving single chain membrane proteins without extra procedures. This data made us one step closer to testing complex membrane protein circuits in cell-free environment.

## Introduction

Cell-free transcription-translation systems have shown to be extremely useful in synthetic biological circuit prototyping [1, 2]. Typical cell-free transcription-translation systems are based on S30 *E. coli* extract [3] and there have been many different versions. The specific version used in this paper referred to as “TX-TL”, has been optimized for prototyping synthetic biocircuits [4, 5]. Unlike other cell-free protein expression systems, including the PURE system, which are based on bacteriophage transcription by supplementing bacteriophage RNA polymerases to the crude cytoplasmic extracts [6], TX-TL has been shown to be suitable for complex biochemical systems [1, 2, 4, 7].

Membrane proteins play important roles in the proper functioning of cells and organisms. They lay the foundation for many biosensors and signal pathways in cells [8]. Some membrane proteins act as ion channels to transport ions across membranes [9]; others make up the essential parts of sensory system which are responsible for cell communication [10], while membrane enzymes catalyze important chemical reactions near membranes [11]. In addition, membrane proteins are the most important drug targets, for example more than half of therapeutics for treatment of various modalities ranging from cancer to cardiovascular diseases target membrane proteins [12].

Membrane proteins are difficult targets as compared to soluble proteins because of the challenges associated with their expression, solubilization, and stabilization. Typical studies of membrane proteins rely on proteins produced from cells and solubilized cell membrane using detergents, liposomes or other membrane-like materials. These approaches may have advantages in terms of protein yield, they are not well-suited for high-throughput selection of protein constructs, since transformation, cell growth, lysis, membrane solubilization and purification are involved for each construct [13]. It is also difficult to use these techniques for biocircuits prototyping, which require prompt function of the newly expressed proteins.

Recently cell free expression of membrane protein have been reported using detergents, liposomes or nanodiscs to solubilize and stabilize the proteins [13–14, 15, 16, 17, 18, 19]. Our data indicated the possibility to integrate membrane proteins directly into future synthetic biocircuits with complex functions. Among all the important membrane proteins, we picked G protein coupled receptors (GPCRs), beta-2(and 1) adrenergic receptors (β2AR/β1AR) as model proteins. With more than 900 members, GPCRs are one of the most important and the largest integral membrane proteins family in human cells and the most important clinical drug targets as they play important roles in many physiological functions and implicated in many diseases [12, 20, 21]. GPCRs all share the same topology – seven transmembrane α-helices, and they are thought to function in monomeric form [22], although there have been studies indicating their dimerization [23, 24]. We used TX-TL platform in combination with nanodisc for expressing β2AR/β1AR and subsequent *in situ* stabilization. Furthermore, biological activity of these proteins was confirmed by binding to its ligands.

## Results and Discussion

We ordered gene synthesis services and then used golden gate assembly [27] to make three different GPCR constructs with three different variants, which share similar backbone, promoter, ribosome binding sites and terminator (Figure 1); Figure 2 lists the difference in coding sequences corresponding to various adrenergic receptor constructs. All three constructs shared the same superfolder GFP (sfGFP) fusion protein topology, which was tagged with 6xHis tag at the C-terminal of the protein. sfGFP is used for monitoring and estimation of target protein expression level during and at the end of TX-TL. Whereas, 6xHis tag was used to detect proteins in Western, to capture the protein for binding assays, and for affinity purification, if necessary.

**Figure 1.**
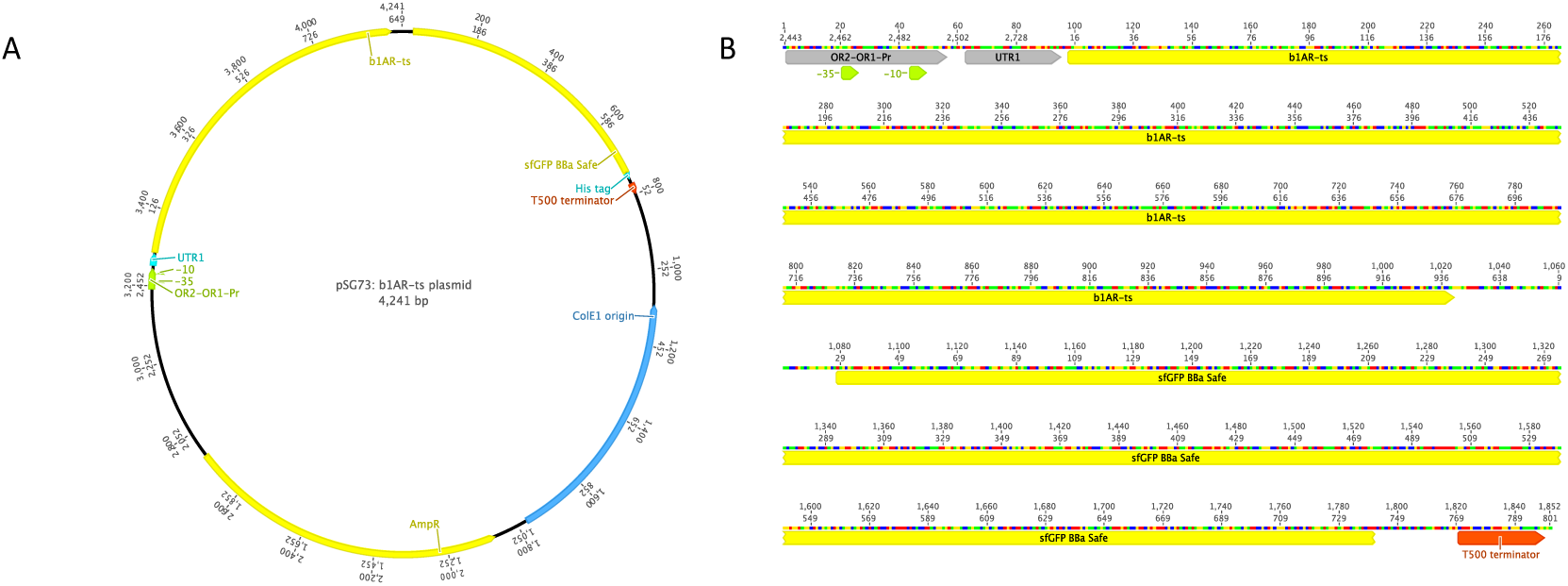
Illustration of the plasmid map and linear DNA of the constructs. pSG73 is used as an example. pSG74 and pSG75 share the same features as pSG73. **A:** Circular plasmid map of pSG73. **B:** Linear DNA version of pSG73.

**Figure 2.**
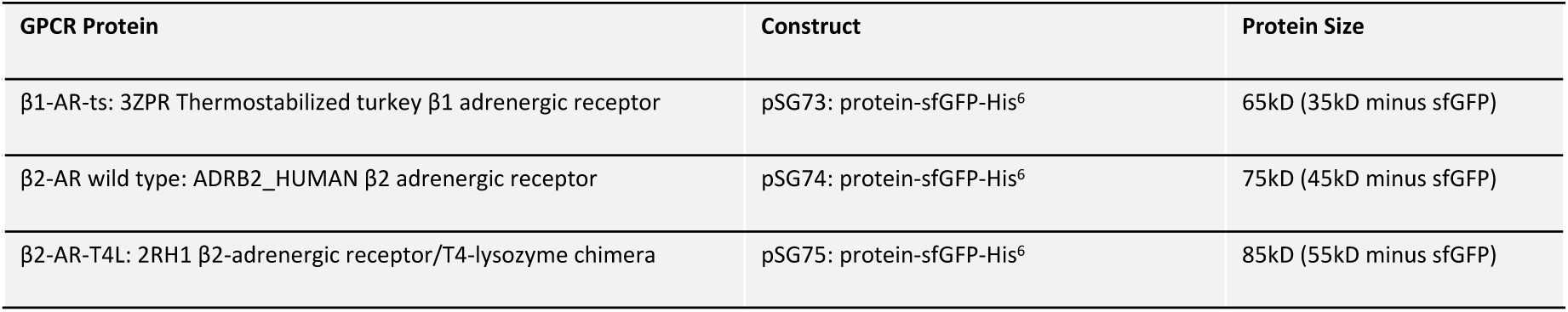
Information of GPCR constructs used in experiments. pSG73 and pSG75 have the corresponding PDB number noted. Protein size of each fusion protein is also listed.

One of the advantages of TX-TL expression platform is that we could use either linear DNA or plasmid DNA for expression in TX-TL [7]. Because of that, we could implement fast construct prototyping in TX-TL by ligating parts together and amplifying the linear DNAs with PCR, avoiding cloning the linear fragments into plasmid. To express these constructs in TX-TL, we only added DNA of these constructs to TX-TL reaction mix. The iteration of each prototyping test significantly faster (overnight) compared to weeks using cloning for cell (*E.coli*) expression.

We first estimated the expression level of pSG73-75 in TX-TL by measuring the fluorescence of sfGFP which was fused at the C-terminal of β2AR or 1AR, using a plate reader at 485 nm (absorbance)/525 nm (emission). All the linear DNA constructs showed a GFP fluorescence signal, indicating successful expression of the fusion proteins in TX-TL (Figure 3A). Different constructs, despite having exactly the same promoter, ribosome binding site, fusion protein framework and DNA concentration, showed different expression levels of sfGFP, especially pSG74. This suggested that the difference in coding sequences could affect transcription and translation level. Another observation was that increasing linear DNA concentration could help increase the fusion protein expression, but this increase was limited by TX-TL resources and/or toxics accumulated in batch mode, as shown elsewhere [28]. Setting up TX-TL reactions in dialysis systems is likely to improve the protein expression level.

**Figure 3.**
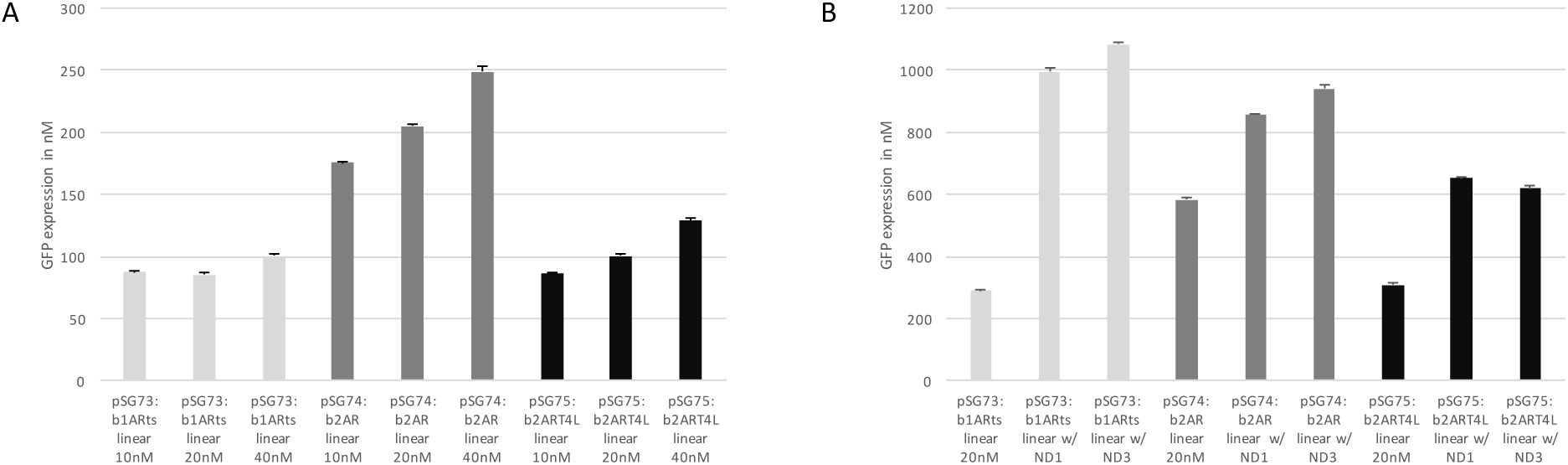
Expression level of pSG73-75 constructs in TXTL measured by sfGFP **A:** End point measurement of GFP expression from 10nM, 20nM and 40nM linear DNA of pSG73, pSG74 and pSG75 in 10μL TX-TL. **B:** End point measurement of GFP expression from 20nM linear DNA of pSG73-75 with or without 24μM nanodiscs in TX-TL. ND1 is MSP1D1-DMPC and ND3 is MSP1E3D1-DMPG. Measurements were done at 29°C in BIOTEK Synergy H1 Hybrid Multi-Mode Microplate Reader using ex485nm/em525nm.

After testing the fusion protein expression level, we explored the possibility of stabilizing hydrophobic membrane protein by providing a membrane mimic into the reaction. Detergent micelle, liposome and nanodisc are commonly used to provide artificial bilayer environment to membrane protein (Ref). We observed that majority of detergents were detrimental to TX-TL reaction comprising membrane protein (data not shown). Conversely, reconstituting protein into liposome resulted into a poor yield of the folded protein, which could be attributed to the liposome’s closed topography. Therefore, we employed nanodisc − a lipid-protein complex, composed of lipids constrained by a membrane scaffold protein (MSP). Nanodisc provides a robust platform with two-dimensional topography for reconstituting TX-TL expressed membrane protein. We chose two of the most commonly used nanodiscs: MSP1D1-DMPC (ND1) ~10 nm diameter and a longer version, MSP1E3D1-DMPG (ND3) ~13 nm in diameter [13].

We repeated the TX-TL expression experiment in the presence/absence of ND1 or ND3. As shown in Figure 3B, the presence of nanodisc improves the TX-TL efficiency, as indicated by enhanced GFP fluorescence. Nanodisc doesn’t have any intrinsic fluorescence, therefore the increased fluorescence should be arising from the fusion protein. GPCR-sfGFP fusion protein precipitated in the TXTL reaction mix without nanodisc. However, in the presence of nanodisc, the supernatant of the reaction mixture exhibited increased GFP fluorescence signal, indicating that nanodiscs likely to help stabilizing fusion membrane proteins in TX-TL, and keep the folded fusion protein in supernatant.

To further confirm that the target fusion proteins were expressed in TX-TL and nanodiscs could help solubilize GPCR-sfGFP fusion proteins, we ran SDS-PAGE of TX-TL samples and western blots with an anti-His6 antibody. First, we ran the whole reaction samples of three different constructs at three different concentrations (Figure 4A).

**Figure 4.**
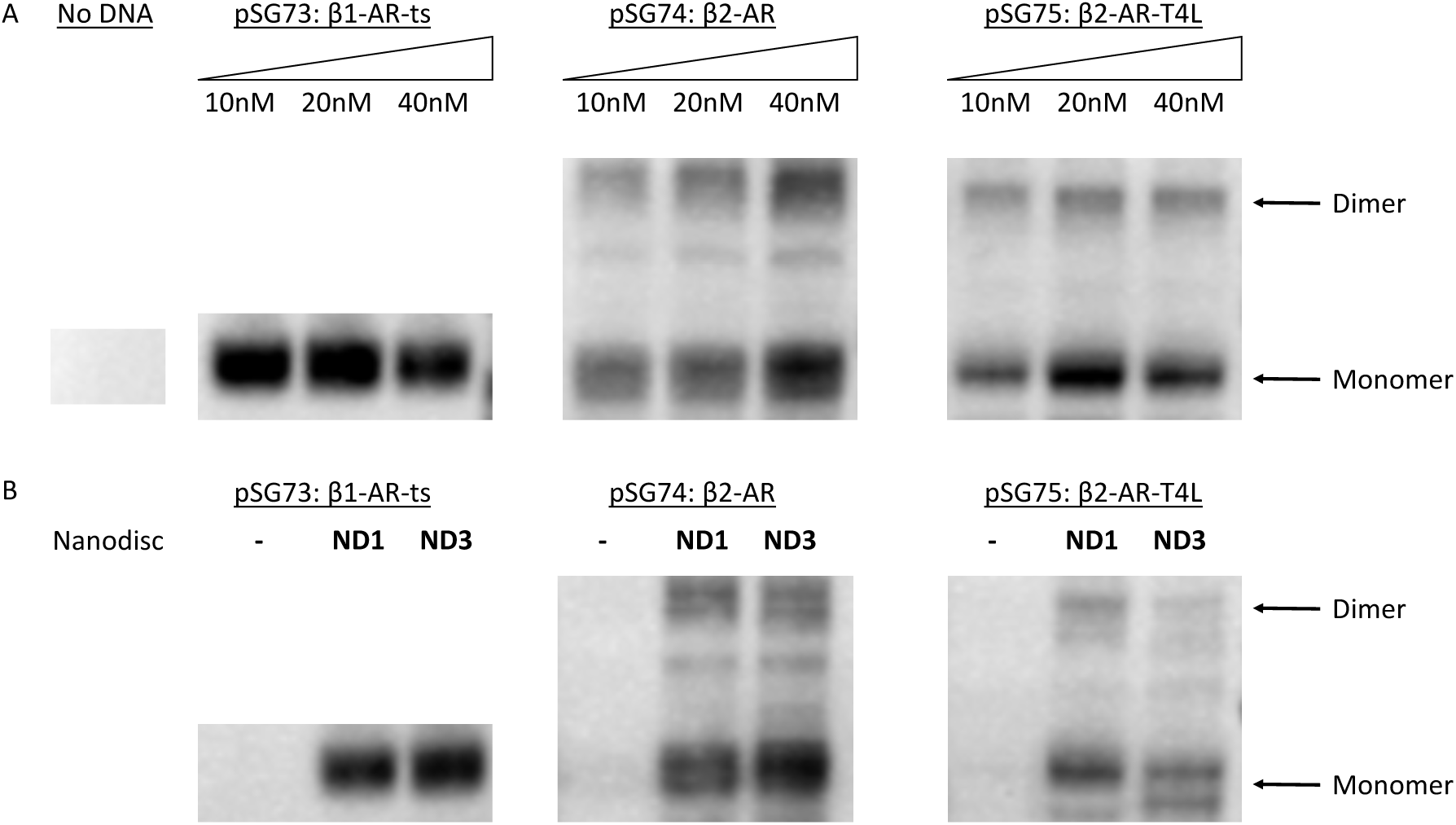
Western blots of TX-TL reactions. **A:** Results of whole reaction TX-TL samples. After measuring the fluorescence of the TX-TL reaction, they were used for western blot using the method in Materials and Methods. TX-TL reaction with no DNA was used as negative control and three different concentrations of three different constructs were run under the same condition. **B:** Results of only supernatant from TX-TL samples. Only soluble samples were run on this blot. Supernatants without any nanodiscs were run at the same condition as ones with either ND1 or ND3.

As illustrated on the blot, samples that had one of the three constructs showed significant detection of His-tagged proteins, protein size was confirmed on the blot using a ladder. In contrast, control sample with without DNA had no His-tagged signal. Additionally, there were not significant differences between different concentrations in pSG73 and pSG75, suggesting mass-transfer nutrients limitation and toxic accumulation in the TX-TL reaction as discussed earlier. Another interesting observation was that pSG73-β1-AR-ts did not show dimerization but pSG74 and pSG75, which were both β2-AR based fusion proteins, showed strong dimer bands on the blot. The presence of GPCR dimerization is consistent with previous reports [24].

We further tested the presence of ND by spinning down the TX-TL reaction mix and took supernatant only to run the western blot. All the insoluble proteins would precipitate out and ended up in the pellet after centrifugation. Only soluble proteins would be in the supernatant and can be detected in the western blot. In Figure 4B, we had three different constructs and each of them had three different experimental conditions: no nanodisc; with ND1: MSP1D1-DMPC; and with ND3: MSP1E3D1-DMPG. All target proteins generated in TX-TL precipitated without nanodiscs and left in the pellet (confirmed by running pellet on western blot, data not shown). On the contrary, when either ND1 or ND3 was added into the reaction, we saw significant bands of target membrane proteins by the Western, suggesting that these hydrophobic membrane proteins became soluble with help from nanodiscs.

So far we have demonstrated that target membrane proteins produced in TXTL reactions can be solubilized and stabilized with in-situ presence of nanodiscs. However, protein association with nanodisc doesn’t insure that these proteins are biologically active. To further test whether these soluble proteins were active, we designed two binding assays: fluorescence-based carazolol binding assay and SPR based norepinephrine binding assay.

Although there is no need to purify the protein for circuit prototyping, purification is required to confirm the binding activity of TX-TL expressed protein to avoid interference caused by *E. coli* endogenous proteins in the TX-TL reaction mix.

We used Ni-NTA affinity chromatography to purify His-tagged target protein. The protocol is described in Materials and Methods.

Fluorescence-based carazolol binding assay uses (S)-carazolol, a derivative of the potent β blocker carazolol with fluorescence properties (ex633nm/em650nm). This ligand can be used as a fluorescence tracker for β2AR binding activity. The detailed experimental setup is described in Materials and Methods. Briefly, purified membrane proteins were incubated with or without carazolol for one hour. These samples were further dialyzed against 100X volume buffer overnight. Subsequently, all the samples were concentrated down to their starting volume and fluorescence signal was measured in plate reader. Green fluorescence (GFP) indicates the amount of fusion proteins in the samples and red fluorescence would represent the amount of carazolol bound to target membrane proteins as an indication of protein activity.

We used two different ND1 (MSP1D1-DMPC) in this assay: 1) HisND uses 6xHis tagged MSP1D1 and was purchased from Cube Biotech; 2) BiotinND uses biotinylated MSP1D1 and was made in house. As seen in Figure 5B, there was not much difference in green fluorescence signal between the samples. However, there is a significant difference between ND1 samples incubated with carazolol. We attribute high carazolol signal on HisND reconstituted β2AR to the presence of empty ND after Ni-NTA affinity chromatography purification.

**Figure 5.**
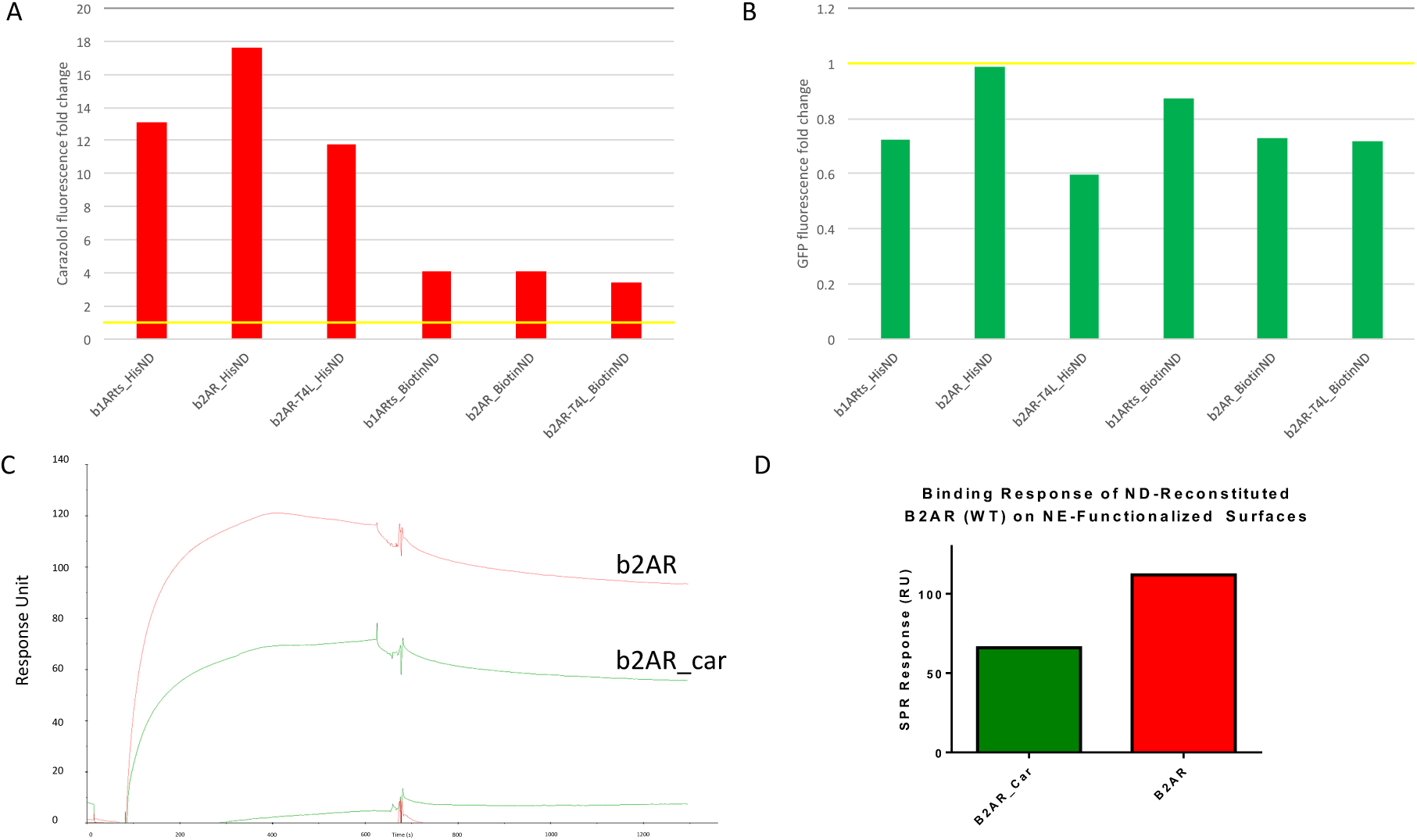
Binding assays of TX-TL made β1 and β2 adrenergic receptor. **A:** Bar chart of carazolol fluorescence fold change. The number was the ratio of red fluorescence signal from samples incubated with carazolol to the ones without carazolol. Detailed experimental method is in Materials and Methods. All samples were dialyzed against buffer without carazolol to remove non-binding carazolol. **B:** Bar chart of GFP fluorescence fold change with the same samples from A. The number was the ratio of green fluorescence signal from samples incubated with carazolol to the ones without carazolol. **C:** Response curves of β2 adrenergic receptor proteins binding to SPR surface coated with norepinephrine. Red curve was response signal from β2-AR sample without carazolol. Green curve was response signal from sample incubated with 1μM carazolol as a control. **D:** Bar chart of the end point of the response curves in C.

One limitation of fluorescence-based assay is that non-specific binding of carazolol to lipids surrounding membrane protein could cause higher signal-to-noise ratio. To overcome this, we developed a SPR-based binding assay using norepinephrine, which is a partial agonist for β2AR. Since norepinephrine shares the same binding pocket with carazolol [31], we tethered norepinephrine to the surface via amide bond between its primary amine and the carboxyl group on the surface. We hypothesized that carazolol could be used as a competitor for β2AR to norepinephrine binding on SPR. As shown in Figure 5C., SPR binding response of β2AR decreased ~60% when it was incubated with 1μM carazolol, indicating specific interaction between β2AR and its binding partners.

To summarize, we have demonstrated that 1) membrane protein can be expressed in TX-TL at analytical scale; 2) presence of nanodisc during TX-TL reaction facilitates folding and solubilization of single chain membrane protein; 3) fluorescence and SPR binding assays were developed to demonstrate specific interaction between small-molecule and nanodisc-stabilized membrane protein.

## Conclusions

In this work, we tested proteins from the GPCR family in our cell-free transcription-translation (TX-TL) system, which would be ideal for synthetic biocircuit prototyping. We expressed β1AR/β2AR in TX-TL with nanodiscs and were able to show that not only these membrane proteins are soluble in TX-TL with nanodiscs, but they were also active. Nanodiscs are co-translationally associated with membrane proteins without extra processing, which enables direct prototyping of a complete biocircuit with TX-TL produced membrane protein(s).

We intend to optimize our binding assays and perform competitive binding experiments to test the stringency of this assay. We envision that GPCR co-expressed with G protein can be expressed by TX-TL to test the biological circuit including signal transduction.

Additionally, histidine kinases, which phosphorylate corresponding response regulators, can activate downstream transcription and translation. We have started tested some hybrid histidine kinases and have seen promising results. Our goal is to prototype a logic biocircuit with membrane enzyme in it, expanding TX-TL platform to broader topics.

## Materials and Methods

### Plasmids and linear DNAs

DNA and oligonucleotides primers were ordered from Integrated DNA Technologies (IDT, Coralville, Iowa). Plasmids in this study were designed in Geneious 8 (Biomatters, Ltd.) and were made using standard golden gate assembly (GGA) protocols. BsaI-HF (R3535S) enzyme used in GGA was purchase from New England Biolabs (NEB). Linear DNAs were made by PCRing protein expression related sequences out of GGA constructs using Phusion Hot Start Flex 2X Master Mix (M0536L) from NEB.

### TX-TL reactions

TX-TL reaction mix was set up according to previous JOVE paper[5]. Briefly, TX-TL extract and buffer were mixed together with calculated linear DNAs or plasmids with or without nanodiscs. Reaction volumes varied from 10μL (initial screening) to 1mL (protein purification and analysis).

### Gel and western blot

Gels used in this work were Bolt 4–12% Bis-Tris Plus Gels from ThermoFisher Scientific. Running buffer was Bolt MES SDS Running Buffer. Gels were run without reducing agents. Protein samples were mixed with LDS sample buffer before loading into gels. iBlot 2 Gel Transfer Device and iBlot Nitrocellulose Regular Stacks were used for transfer proteins from gel to membrane. Membrane was then transferred to iBind device and incubated with Penta-His HRP Conjugate in 1:500 dilutions. Blots were detected using SuperSignal Chemiluminescent HRP Substrates from ThermoFisher Scientific.

### Protein purification

TX-TL reaction mix was first spun @14,000g for 10min at 4°C. Supernatant was then transferred to buffer-equilibrated HisPur Ni-NTA Spin Purification column (ThermoFisher Scientific) and incubated with shaking for 1h. Then spun down the flow through @2000g for 2min and washed with nanodisc buffer (20 mM Tris pH 7.4, 0.1 M NaCl) added with 20mM imidazole three times. Elution was done by adding elution buffer (20 mM Tris pH 7.4, 0.1 M NaCl, 250mM imidazole) for 3 times 2 column volume. Proteins were then concentrated using Amicon Ultra Centrifugal Filter Units (Millipore) with Ultracel-30 membrane.

### Fluorescence-based carazolol binding assay

Carazolol was purchased from Abcam (S)-Carazolol Fluorescent ligand (Red) ab118171. Each purified protein was first divided into two equal volume samples. One was added 100nM carazolol (dissolved in water) and the other one was added the same volume of water. They were incubated at 4°C for 1h before transferred to mini 10k MW D-Tube Dialyzers (Millipore) and dialyzed against 100x volume of nanodisc buffer for overnight. Then dialyzed samples were concentrated using Amicon Ultra Centrifugal Filter Units (Millipore) with Ultracel-30 membrane to starting volume. Samples were then put into BIOTEK Synergy H1 Hybrid Multi-Mode Microplate Reader and measured for GFP fluorescence (ex485nm/em525nm) with gain 61 or gain 100 or Red fluorescence (ex633nm/650nm) with optimal gain. GFP fluorescence was converted to nM using calibration data from purified GFP protein.

### Surface Plasmon Resonance (SPR) based norepinephrine binding assay

GE Biacore T-200 SPR system was used for SPR experiment. Gold plated chip was first immobilized with norepinephrine and then washed away extra chemical. There are four channels on one chip. Two were used as experimental channels and the remaining two were used as negative controls to provide background response from buffer. One sample was incubated with 1μM carazolol for 1h at to test binding specificity and the other sample was incubated with same volume of water. Samples were then loaded and flowed through corresponding experimental channels and response curves were recorded.

## Acknowledgments

We would like to thank Hao Chen for receptor plasmid design. Marie Wright and Robert Kurzeja for nanodisc expression, and Amgen Inc. for financial support.

